# The effect of periodic disturbances and carrying capacity on the significance of selection and drift in complex bacterial communities

**DOI:** 10.1101/2021.07.18.452870

**Authors:** Madeleine Stenshorne Gundersen, Ian Arthur Morelan, Tom Andersen, Ingrid Bakke, Olav Vadstein

**Author notes:** Address correspondence to Madeleine Gundersen.

## Abstract

Understanding how periodical disturbances affect the community assembly processes is vital for predicting temporal dynamics in microbial communities. The effect of dilutions as disturbances are poorly understood. We used a marine bacterial community to investigate the effect of disturbance (+/−) and carrying capacity (high/low) over 50 days in a dispersal-limited 2×2 factorial crossover study in triplicates. The community’s disturbance regime was crossed halfway. We modelled the rate of change in community composition between replicates and used this rate to quantify selection and ecological drift. The disturbed communities increased in Bray-Curtis similarity with 0.011±0.0045 (Period 1) and 0.0092±0.0080 day^−1^ (Period 2), indicating that selection dominated community assembly. The undisturbed communities decreased in similarity at a rate of −0.015±0.0038 day^−1^ in Period 1 and were stable in Period 2 at 0.00050±0.0040 day^−1^, suggesting drift structured community assembly. Interestingly, carrying capacity had minor effects on community dynamics. This study is the first to show that stochastic effects are suppressed by periodical disturbances resulting in exponential growth periods due to density-independent biomass loss and resource input. The increased contribution of selection as a response to disturbances implies that ecosystem prediction is achievable.

## Introduction

Understanding how ecological assembly processes create temporal patterns in community composition is a major goal in community ecology (1). After decades of debating whether community assembly follows neutral (2) or niche theory (3), it is now generally accepted that both stochastic and deterministic processes are important for community assembly (1,4,5).

Four high-order processes structure community assembly. These are selection, ecological drift, dispersion and diversification (4,5). These four processes have a varying degree of stochasticity and determinism. Selection is deterministic and based on differences in the fitness between populations. This process includes environmental filtering and biological interactions, such as competition and mutualisms. Drift is an entirely stochastic process that arises because there is a non-zero probability that an individual dies before it reproduces (6). The outcome of drift is a change in the relative abundance of populations and can lead to local extinction if the abundance is low. Dispersion and diversification are two processes that are both stochastic and deterministic. Dispersion refers to an individual’s movement from the regional to the local species pool, whereas diversification is the evolution of new strains (4). The relative contribution of these four processes on community assembly can vary between sites and changes over time (7,8).

Only experiments with high temporal resolution can evaluate the relative importance of these community assembly processes (9,10). During the last decade, studies using high temporal resolution sampling approaches have pointed to stochastic processes as being more important and selection as less important than previously assumed. This observation has been done in habitats such as bioreactors (11,12), soil (13,14), and wastewater treatment plants (15). This increased awareness of stochasticity emphasises the need for more knowledge on temporal variation in the assembly processes.

A primary motivation for studying microbial community assembly is to understand the communities’ responses to drivers affecting the high-order assembly processes in order to be able to forecast and manage them (10,16). Such control is vital in, for example, treating dysfunctional human gut microbiomes (17), ensuring stability during biological wastewater treatment (18) and providing an optimal microbial environment for fish in aquaculture (19).

Microbial communities often experience disturbances. Disturbances usually involve alterations in the available resources or the biomass concentration in the given environment. To predict the consequence of disturbances on the dynamics of microbial communities, it is essential to understand how the disturbance influences the four assembly processes’ relative contributions (7,10). Some studies have shown that disturbances affect the relative contribution of the assembly processes (e.g. 11,16–21), but conclusions vary depending on the disturbance type and the ecosystem studied. Disturbances that increase resource availability have been found to enhance the contribution of stochastic processes in dispersal limited communities (11,23). This enhancement is likely due to weakened competition and strengthened priority effects (i.e., the effect of colonisation history) due to the increase in resources (7,11,23,26). Conclusions are the opposite for disturbances that result in biomass loss where the contribution of deterministic processes appears to increase (24,25). How community assembly is affected by disturbances that combine resource increase and biomass loss is not known.

The maximum biomass an ecosystem can sustain is controlled by the carrying capacity. With regards to community assembly, carrying capacity can affect drift. Lower carrying capacities support lower biomasses, and as drift is density-dependent, more populations are vulnerable to extinction (6). To our knowledge, no one has investigated how carrying capacity affects community assembly processes.

Microbial microcosms are excellent systems to study the effect of disturbances and carrying capacity on the temporal changes in community assembly. This is due to the short generation time of microorganisms, the potential for high experimental control and the possibility to include many experimental units (27). In microcosms, one can eliminate dispersal, and if community composition is monitored by clustering 16s-rDNA sequences at a 97% similarity level, speciation is negligible (28). Consequently, selection and drift are the only assembly processes shaping the bacterial communities (27).

Selection and drift can be quantified by investigating the similarity in community composition between biological replicates in systems without dispersal and speciation (Figure 1). This approach assumes that if the selection is homogeneous (i.e., there is one stable equilibrium per condition), communities of replicate microcosms should over time become more similar if selection dominates and less similar if drift predominates. Moreover, if selection dominates, one expects the variation in community composition between replicate microcosms to decrease because the communities become more similar over time. Conversely, if drift dominates community assembly, replicates are expected to become less similar, and the variation in compositional similarity will increase with time.

**Figure 1:**
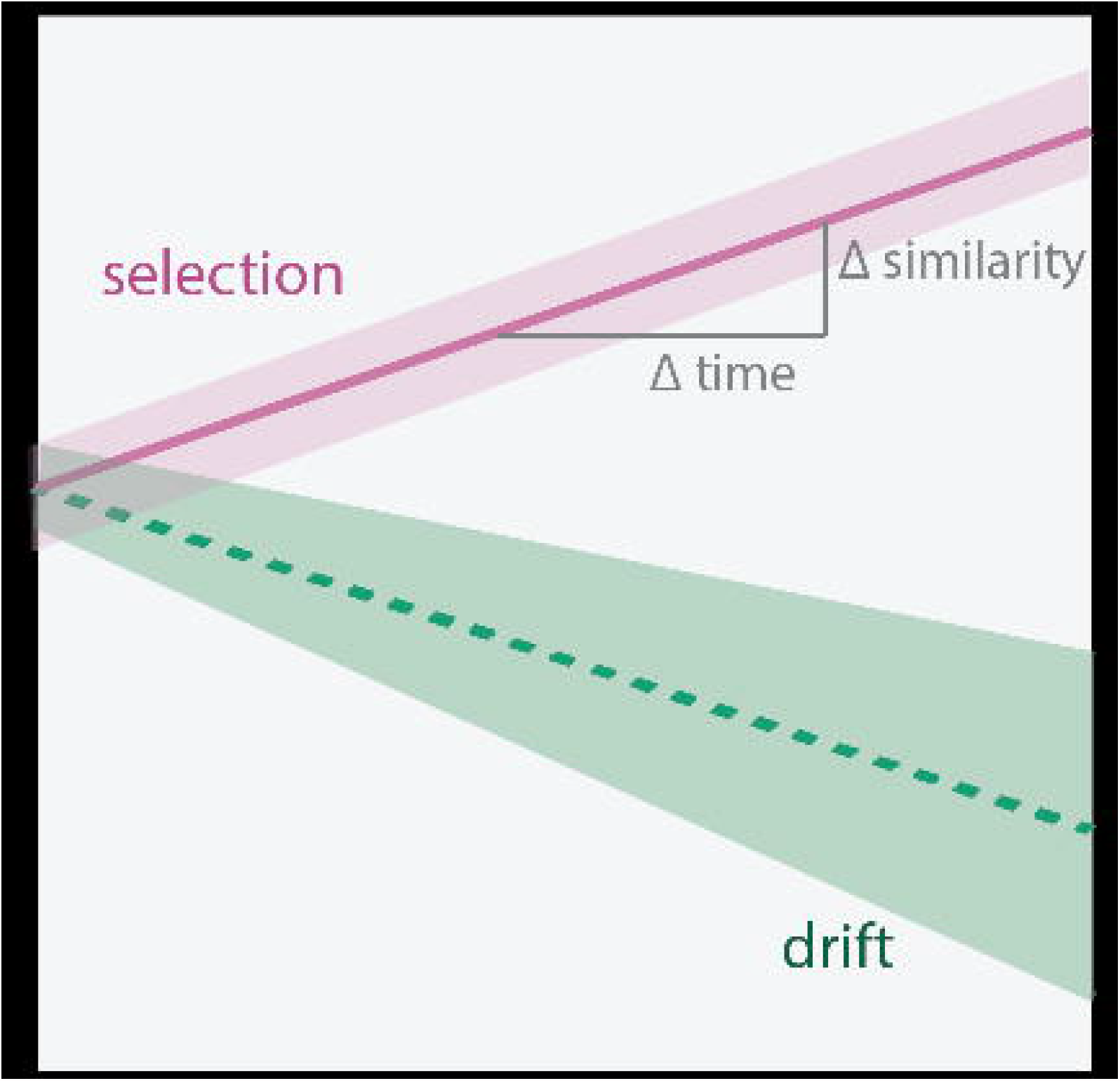
A conceptual schematic of the temporal changes in community similarity between replicates if drift or selection dominates the community assembly. If selection dominates, the similarity between replicates increases over time, and the variance decrease or be stable. However, if drift dominates, replicates should become less similar over time, and the variance should increase.

In the present study, we aimed at disentangling the effects of disturbance introduced as periodic dilutions (undisturbed *versus* disturbed microcosms) and carrying capacity (high *versus* low) on succession and the relative importance of the assembly processes selection and drift. We investigated the effect of these factors using a 2×2 factorial crossover experimental design with three replicate microbial microcosms for each condition over 50 days. The marine microbial communities were cultivated either in chemostats or with semi-continuous cultivation with a 50-fold dilution every second day. The dilution functioned as a combined disturbance as it both reduced the community size and increased the specific resource supply. We quantified selection and drift using the approach described above, which allowed us to gain insight into the effect of disturbance and carrying capacity on selection and drift.

## Materials and methods

### Experimental design and sampling scheme

A marine bacterial community collected from 70 m depth in the Trondheimsfjord, Norway (March 2018) was sand-filtered and used to inoculate twelve microcosms (500 mL, GLS 80® stirred reactor, Duran, Germany) in a 2⨯2 factorial crossover design (Figure 2). Each microcosm contained 250 mL culture that was stirred continuously (MIX 6, 2mag AG, Germany), supplied with 0.2 µm filtered (Millipore) hydrated air, and kept at 15°C. The communities were cultivated in f/2 medium (29) with either 0.33 (low carrying capacity, L) or 5×0.33=1.67 mg/L (high carrying capacity, H) of yeast extract, peptone and tryptone. The inorganic nutrients in the f/2 media were 50-fold diluted compared to the original recipe. The medium was either supplied continuously at a dilution rate of 1 day^−1^ (Watson Marlow 520S peristaltic pump) or pulsed by a 1:50 dilution every second day equivalent to a continuous dilution rate of approximately 2 day^−1^ (Figure 2a and b). We define the pulsed communities as disturbed (D) and those continuously supplied with medium as undisturbed (U). This disturbance regime was crossed after 28 days so that previously disturbed microcosms were undisturbed the last 22 days (DU) and vice versa (UD). The cultivation regimes are abbreviated as UDH, UDL, DUH, and DUL (Figure 2c). Each cultivation regime was run in triplicates. The bacterial communities were sampled by filtering approximately 30 mL of culture through a 0.2 µm filter to a total of 206 samples (2 inoculum and 17 time-points x 4 regimes x 3 replicates) which were stored at −20°C until further processing. Sampling of the disturbed communities was done right before the dilution.

**Figure 2:**
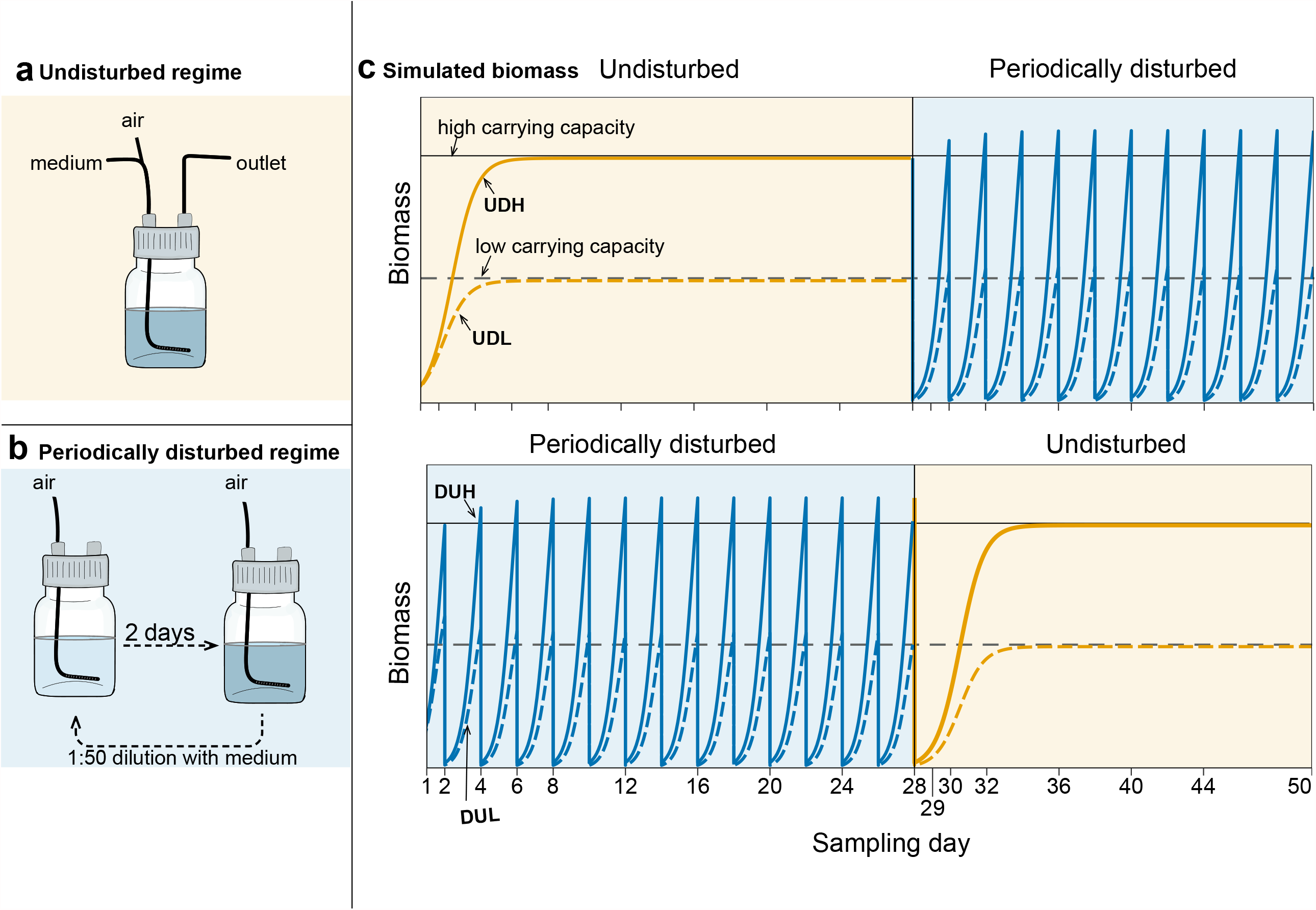
Schematic overview of the experimental design. a) Undisturbed communities (U) received medium continuously at a dilution rate of 1 day^−1^. b) Disturbed communities (D) were 50-fold diluted with medium every second day. The dilution acted as a disturbance because the community’s biomass was reduced substantially, and resources were introduced as a large pulse. c) Simulated logistic growth of the microbial communities’ biomass given the disturbance regime and carrying capacity (parameters: µ = 2.5 day^−1^, continuous dilution = 1 day^−1^ or semi-continuous 1:50 dilution every 2nd day). High carrying capacity is indicated as a solid black line, whereas low carrying capacity is presented as a dashed black line. When the communities are undisturbed, the biomass is expected to be at or near carrying capacity. The disturbance by dilution lowered the community’s biomass by a factor of 50, bringing the community considerably below the carrying capacity, resulting in close to exponential growth between dilutions. For each experimental condition, triplicate microcosms were operated over 50 days, and the disturbance regime was switched after 28 days. The groups are abbreviated as UDH, UDL, DUH and DUL, in which the first letter indicates the disturbance regime in period 1 (day 1-28), the second the disturbance regime in period 2 (day 29-50) and the third letter the carrying capacity of the media (high or low).

### Extraction of Bacterial DNA and 16S-rRNA amplicon sequencing

Bacterial community DNA was extracted with the Qiagen DNeasy PowerSoil DNA extraction kit. The V3-V4 region of the bacterial 16S-rRNA gene was amplified using the broad-coverage primers with Illumina MiSeq adapter sequences Ill338F (5’-TCG-TCG-GCA-GCG-TCA-GAT-GTG-TAT-AAG-AGA-CAG-NNN-NCC-TAC-GGG-WGG-CAG-CAG-3’) and Ill805R (5’-GTC-TCG-TGG-GCT-CGG-AGA-TGT-GTA-TAA-GAG-ACA-GNN-NNG-ACT-ACN-GG-GTA-TCT-AAK-CC-3’). The reactions were run for 28 cycles (98°C 15 s, 55°C 20 s, 72°C 20 s) with 0.3 µM of each primer, 0.25 mM of each dNTP, 1 mM of MgCl_2_, 2 µM of 5x Phusion buffer HF, 0.015 units/µL of Phusion Hot Start II DNA polymerase, 1 µL of DNA template and dH_2_0 to a total volume of 25 µL. The amplicon library was prepared as described previously (30). In brief, we used the SequalPrep Normalisation plate (96) kit (Invitrogen) to normalise and purify PCR products and the Illumina Nextera XT Index kits (FC-131-2001 and FC-131-2004) for amplicon indexing. The amplicon library was sequenced with V3 reagents by 300 bp paired-end reads on two MiSeq Illumina runs at the Norwegian Sequencing Centre. Illumina sequencing data are deposited at the European Nucleotide Achieve (accession number X – to be uploaded to ENA).

### Processing of sequence data

We used the USEARCH pipeline (v11) to process the Illumina sequence data (31). Briefly, using the command fastq_mergepairs, paired ends were merged, and primer sequences and reads shorter than 400 bp were removed. The data was quality filtered using the command fastq_filter with an expected error parameter of 1, and singletons were removed. We used the UPARSE-OTU algorithm to remove chimaeras and cluster OTUs at the 97% similarity level (32). Taxonomy was assigned to the OTUs using the Sintax command with the RDP reference dataset (RPD training data set version16) at an 80% confidence threshold (33,34).

### Analysis of diversity and differential abundance testing

The resulting OTU-table was further analysed in R (version 3.6.1) (35). All R-code is provided at https://github.com/madeleine-gundersen/disturcance-cc-assembly. We first evaluated the sequencing effort with the function rarecurve() in the vegan package (version 2.5-6) (36). Then the data were normalised by averaging 1000 rarefied datasets created by randomly sub-sampling 10 000 reads without replacement using phyloseq_mult_raref() from the package metagMisc (version 0.0.4) (https://github.com/vmikk/metagMisc/).

Alpha diversity was estimated as Hill diversity of order 0 to 2 (37) with the function renyi() in vegan. Bray-Curtis and Sørensen similarity indices were used to quantify beta diversity (38). The variance in beta-diversity was ordinated with Principal Coordinate analysis (PCoA) (39). Permutational multivariate analysis of variance (PERMANOVA) was used to test if sample groups significantly differed in community composition. The effect size of variables was evaluated with the R^2^-value estimated with PERMANOVA (40). To determine which OTUs increased in abundance due to the disturbance regimes, we performed a differential abundance test with DeSeq2 (41). We used the non-normalised OTU-table as input to the DeSeq2 analysis. Only samples from the last two weeks of the cultivation periods were included in the analysis as PCoA ordinations indicated that the communities had stabilised. First, the abundance data were normalised using the median ratio method. DeSeq2 was then run with the Wald significance test assuming the negative binomial distribution. All p-values were FDR corrected.

### Estimation of selection and drift on community composition

We developed a new approach to quantify the contribution of selection and drift during succession in highly controlled experimental settings where dispersal and speciation can be negligible (Figure 1). Our approach is based on a three-step analytical process. First, the similarity in community composition between replicate pairs is calculated at each sampling day. Then the change in similarity is regressed again time. Finally, the slope of the temporal change in similarity is used to quantify selection and drift. Selection will result in communities becoming more similar with time, resulting in a positive or neutral slope. In contrast, drift causes communities to become less similar over time, manifested as negative slopes. In addition to the slope, the variation in similarity measurements can strengthen the conclusions as selection should decrease variation. In contrast, drift should increase the variation.

We calculated pair-wise community similarities between replicate microcosms at each sampling day, using Bray-Curtis and Sørensen similarity indices. In the following, we will use the term “replicate similarity” for this metric. We used a hierarchical Bayesian model approach to estimate the rate of change in the replicate similarity. We chose a Bayesian approach as it has the advantage of accounting for this dataset’s hierarchical dependencies, few observations per time point and the observed heteroscedastic variance (26,42).

We fitted hierarchical linear Bayesian models with replicate similarity as the dependent variable using the brms package (version 2.11.1) (43), which is a user-friendly front-end for the Stan system for Bayesian computing (44). All models had a random intercept term for the three similarity comparisons (+(1|comparison) in each time and regime combination. We modelled the replicate similarity by a normal distribution with fixed effects on both mean (µ) and standard deviation (σ) using default priors. Fixed effects included 3-way interactions between time, disturbance, and carrying capacity for the mean model, whereas the standard deviation model only had interactions between time and disturbance. We mean-centred the time variable to reduce correlations between fixed effect estimates. MCMC simulations with brms were run on 4 chains with 4000 samples each (2000 for warm-up), giving 8000 posterior samples. To reduce the number of divergent transitions in the MCMC sampling, we increased the value of the adapt_delta parameter to 0.99 (default is 0.95). We fitted several models, compared their predictive densities, and selected the model structure with the highest predictability for the temporal development of similarities between replicates. An overview of all estimated models and model selection process is given in the Supplementary material.

We used the package tidybayes (version 2.0.3, http://mjskay.github.io/tidybayes/) to extract posterior samples and the stat_lineribbon() aesthetic from ggplot2 (45) to visualise fixed effect means and credible intervals of model predictions. As explained above, we interpreted community assembly as being dominated by selection if the time effect on the mean of replicate similarity was non-negative (i.e., µ day^−1^ ≥ 0), and the standard deviation slope was non-positive (σ day^−1^ < 0) (Figure 1). Conversely, we interpreted a negative slope for the mean and a positive slope for the standard deviation as a community assembly dominated by drift (i.e., µ day^−1^ < 0, σ day^−1^ > 0). In cases where the fit met neither of these criteria, we defined the community assembly as a mix of selection and drift.

## Results

To study the effect of the periodical disturbance and carrying capacity on community succession and the assembly processes, we cultured marine microbial communities under the DUH, DUL, UDH, and UDL cultivation regimes and characterised their temporal dynamics using 16S-rDNA amplicon sequencing. The dataset contained a total of 12 945 783 sequence reads with a mean of 63 460 reads (± 31 411 SD) per sample. The dataset was normalised to 10 000 reads per sample (**Error! Reference source not found**.). The Hill alpha diversity of order 0, 1 and 2 of the normalised dataset correlated well with the non-normalised dataset (p<0.05). The slopes of linear regressions between the alpha diversities of these datasets were close to one, indicating that the normalised- emulated the non-normalised dataset (**Error! Reference source not found**.). The samples from the first sampling day were removed from the dataset because the richness dropped 43% from day 1 to 2 (**Error! Reference source not found**.). This reduction was probably an adaption of the original seawater community to the culture conditions. During the rest of the experiment, the richness was relatively stable, and a total of 739 OTUs were observed for the normalised OTU table (**Error! Reference source not found**.).

### The disturbance regime drove succession

The community succession differed between the cultivation regimes, as indicated by PCoA ordinations based on both Bray-Curtis (Figure 3: day 16-28, 36-50, **Error! Reference source not found**.: day 2-50) and Sørensen dissimilarities (**Error! Reference source not found**.: day 16-28, 36-50). Disturbance accounted for over 44 and 50% of the variation in Bray-Curtis dissimilarities at the end of Period 1 and 2, respectively (p<0.001, PERMANOVA). Carrying capacity accounted for only 6 (p=0.16) and 11% of the variation (p = 0.04) for the two periods. A fascinating observation was that switching the disturbance regime resulted in a reversal of the community succession from the undisturbed ordination space to the disturbed one, and vice versa (Figure 3, **Error! Reference source not found**.). These ordinations indicated that disturbance was the main contributor to the succession and that carrying capacity had less effect.

**Figure 3:**
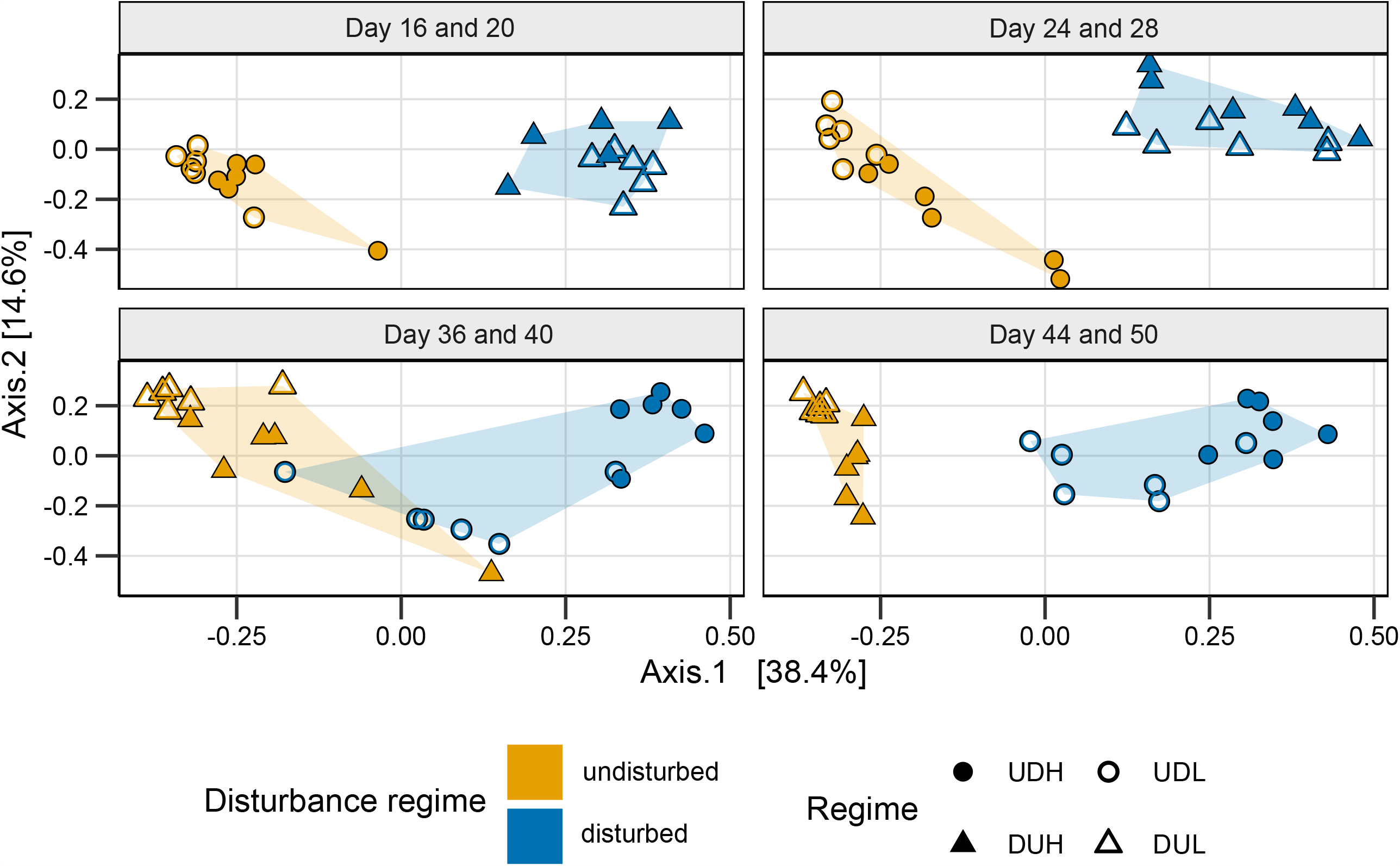
PCoA ordination based on Bray-Curtis dissimilarity for the bacterial communities at the end of Period 1 (day 16-28) and Period 2 (36-50). The single ordination for these samples was split by sampling-week to highlight the succession based on the disturbance regime. UD (circles) were undisturbed the first 28 days and disturbed the last 22 days, while DU (triangles) were disturbed in the first period and undisturbed in the second. H (filled) and L (empty) indicates high and low carrying capacity, respectively. Colours represent the disturbance regime at sampling, and the shaded area the spread of samples with similar disturbance regime.

Comparing the replicate similarity at the start and the end of each cultivation period indicated that the communities became more similar during disturbance than when undisturbed (Figure 4a). For the disturbed communities, the Bray-Curtis similarity increased by 138% during Period 1 (DU) and 46% during Period 2 (UD). In contrast, the undisturbed communities decreased in similarity by 47% during Period 1 (UD) and increased by only 3.9% during Period 2 (DU, Figure 4a). We investigated the replicate similarity change over time to determine whether selection or drift structured these successional patterns.

**Figure 4:**
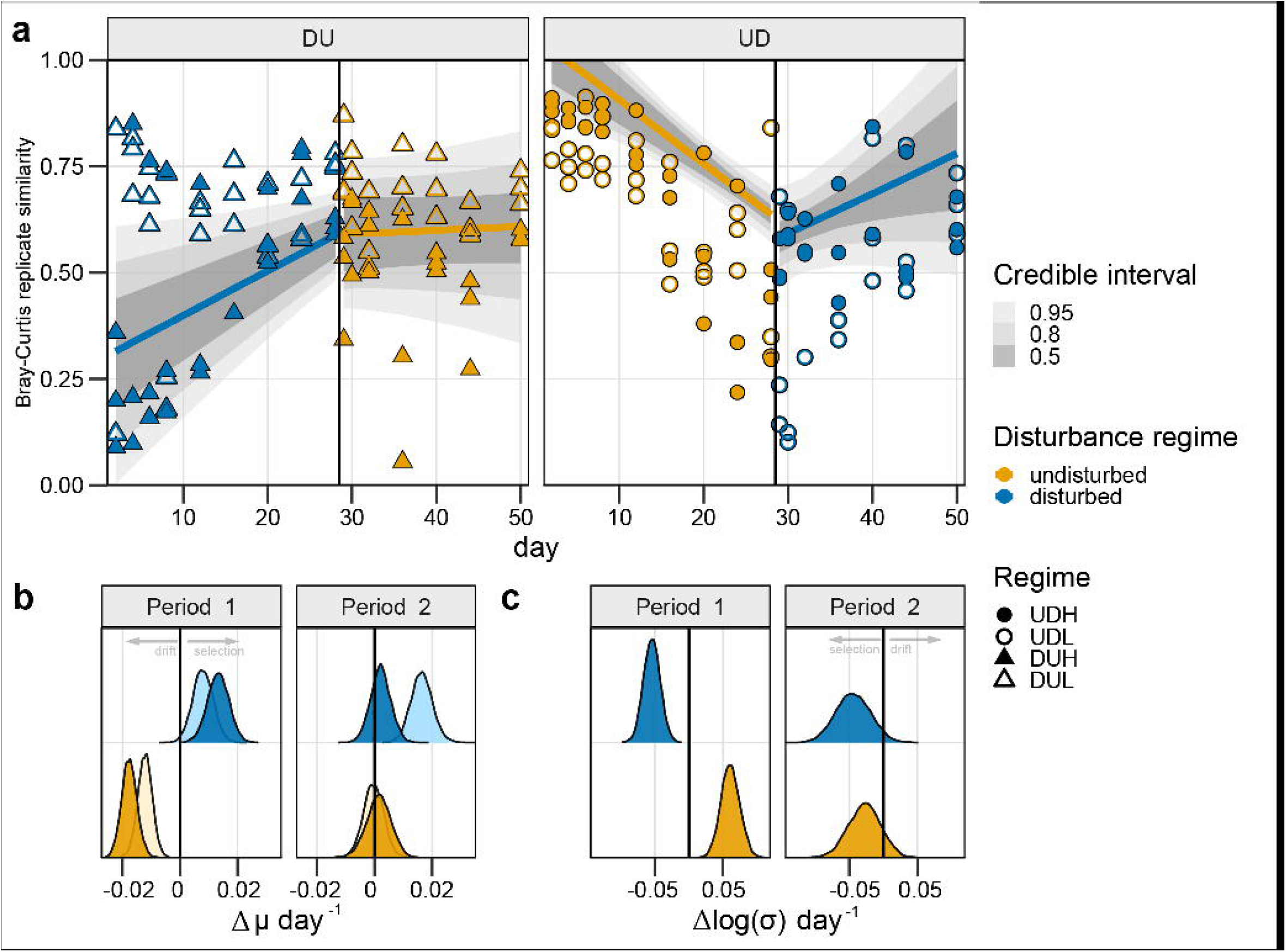
The Bray-Curtis based models and coefficient estimates for the change in the similarity between replicates over time. **a)** The similarity between replicate communities as a function of time. The models for the replicate similarity change over time are presented as lines with the 0.5, 0,8 and 0.95 credible intervals around it. The observed data used as the response variable in the models are presented as points. UD (circles) were undisturbed the first 28 days and disturbed the last 22 days, while DU (triangles) were disturbed in the first period and undisturbed in the second. H (filled) and L (empty) indicates high and low carrying capacity, respectively. Colours represent the disturbance regime at sampling. **b)** The posterior distributions of the expected replicate similarity (µ) change per day given the interaction between time, the disturbance regime and carrying capacity. The distribution reflects all 8000 estimated replicate similarity changes that would give the observed data. Light and dark colour indicate low and high carrying capacity, respectively. The colour indicates the disturbance regime at sampling. **c)** The posterior distributions for the change in standard deviation per day given the interaction between time and disturbance regime. The distribution reflects all 8000 estimated standard deviation changes per day that would give the observed data. The colour indicates the disturbance regime at sampling.

### Selection dominated during disturbance

We used a Bayesian hierarchical model approach to estimate the replicate similarity change over time, and based on this, we examined whether selection or drift dominated the successions. The deterministic process selection should result in communities increasing in replicate similarity over time. This increased similarity will also result in a decrease in variation between similarity measurements. In contrast, the random process drift would reduce the replicate similarity and increase the variation over time (see Figure 1 and Materials and method for more information).

For the disturbed communities in Period 1 (DU), the posterior-distributions of the model parameters revealed that the replicate similarity increased with 0.011±0.0045 day^−1^ (mean±SD), whereas the standard deviation decreased 0.054±0.010 day^−1^ (Figure 4b and c). This increased replicate similarity and decreased standard deviation over time indicate that selection was the dominating assembly process (Figure 5a). Moreover, we observed the same trends for the disturbed communities in Period 2 (UD) with an increase in replicate similarity of 0.0092±0.0080 day^−1^ and a decrease in the standard deviation of −0.044±0.025 day^−1^ (Figure 4b and c). This coherent observation strengthens the conclusion that selection dominated community assembly during disturbances.

**Figure 5:**
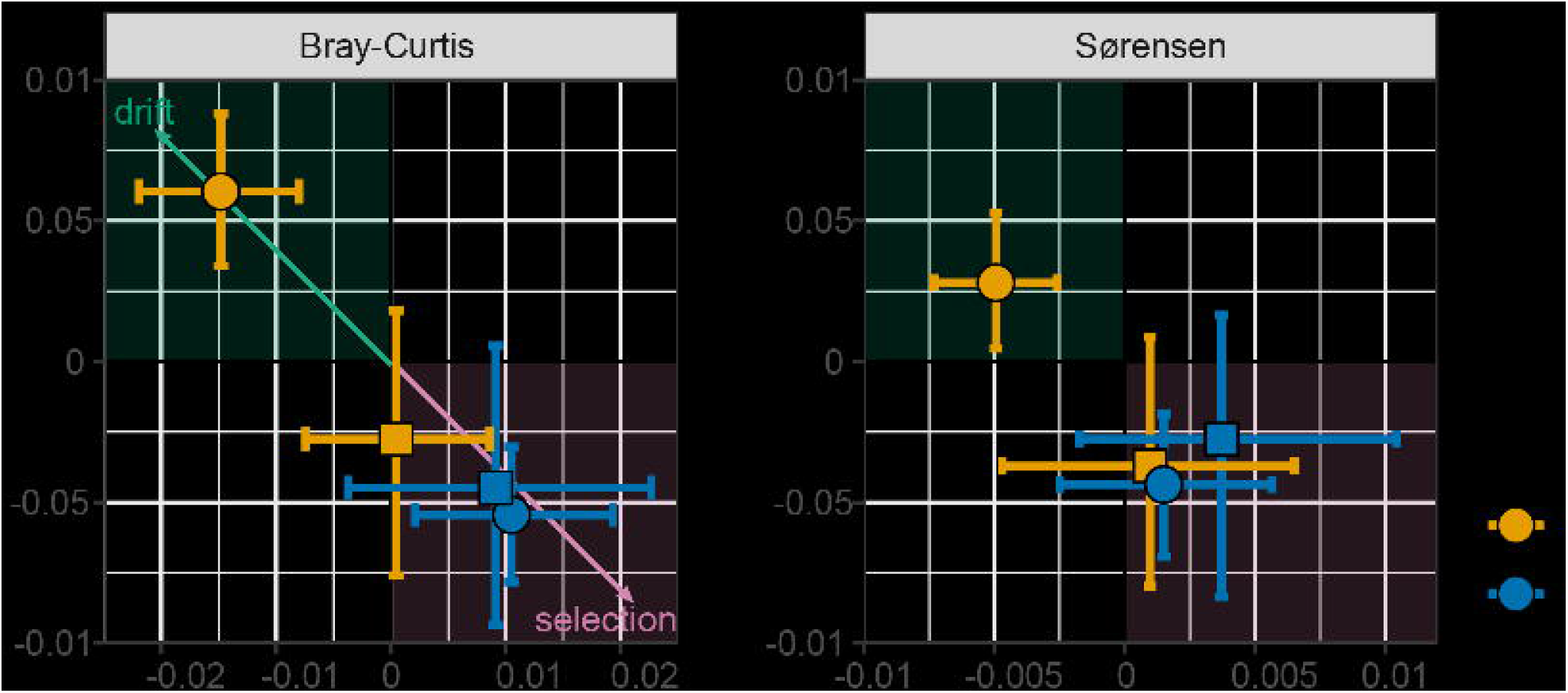
The mean change in replicate similarity (µ) over time and mean change in standard deviation (σ) (points) and the accompanying 95% credible intervals for each estimate, as inferred using the Bayesian hierarchical model approach, based on a) Bray-Curtis and b) Sørensen based models. The green area indicates the coordinate space where drift dominates, while the pink areas indicate where selection is dominating. Point colour indicates the disturbance regime and shapes the cultivation period.

The modelling results were different for the undisturbed communities. During Period 1 (UD), the replicate similarity rate decreased by −0.015± 0.0038 day^−1^ and had a temporal increase in variation of 0.061±0.014 day^−1^ (Figure 4b and c), indicating that drift dominated the community assembly (Figure 5a). For the communities that switched from a disturbed to an undisturbed regime (DU) in Period 2 the dominating assembly process was less obvious. The replicate similarity rate was relatively stable with a mean of 0.00050±0.00400 day^−1^ and a decrease in the standard deviation of −0.028±0.024 (Figure 4b and c). These values categorise the assembly as selection (Figure 5). However, comparing the replicate similarity rate of the communities from Period 1 to the one in Period 2 shows that the rate decreased substantially. The average similarity rate transitioned from the selection-coordinate space towards the one where drift dominates (Figure 5a).

The results were similar for models based on the Sørensen similarity, with an overall increase in replicate similarity over time for the disturbed regimes of 0.0015±0.0021 day^−1^ in Period 1 (DU) and 0.0041±0.0040 day^−1^in Period 2 (UD), and a decrease in the standard deviation of the replicate similarity over time (Figure 5b, **Error! Reference source not found**.). For the undisturbed communities, there was a slight temporal decrease in Sørensen replicate similarity at a rate of −0.0049±0.0012 day^−1^ in Period 1 (UD), whereas in Period 2 there was an insignificant change in replicate similarity (0.00096±0.0029 day^−1^; DU). These results supported the findings based on the Bray-Curtis similarity; drift dominated assembly for the undisturbed communities, whereas selection dominated when the communities were disturbed.

### *Gammaproteobacteria* increased in relative abundance during disturbance

The PCoA ordination and the replicate similarity models showed that the disturbance regime impacted the assembly processes. We performed a DeSeq2 differential analysis to elucidate which OTUs had significantly different abundances between the disturbed and undisturbed regime. This analysis revealed that 107 of the 535 OTUs contributed significantly (p<0.05) to differences in community composition between the disturbed and the undisturbed regimes. These OTUs were grouped at the genus level (Figure 6). Interestingly, around 60% of these groups included only one OTU. For genera with more OTUs affected, the general trend was that the OTUs responded similarly to the disturbance regime (i.e. either positive or negative fold change in relative abundance). For example, all 13 OTUs classified as *Colwellia* and all 5 OTUs classified as *Vibrio* had higher abundance during disturbance. However, this was not the case for all the groups. For example, of the 21 OTUs classified to *Rhodobacteraceae*, 6 were in higher abundances during the disturbed periods, whereas 15 were more abundant during undisturbed periods. Thus, some genera’s OTU abundances appeared to respond to the disturbance regime coherently, whereas others did not. Of the 107 OTUs significantly affected by the disturbance regime, 72% had increased abundances when the environment was disturbed. Especially noteworthy was the *Gammaproteobacteria*, where 94% of the OTUs significantly affected by the disturbance regime had higher abundances during disturbance with up to an 11.2 fold-change.

**Figure 6:**
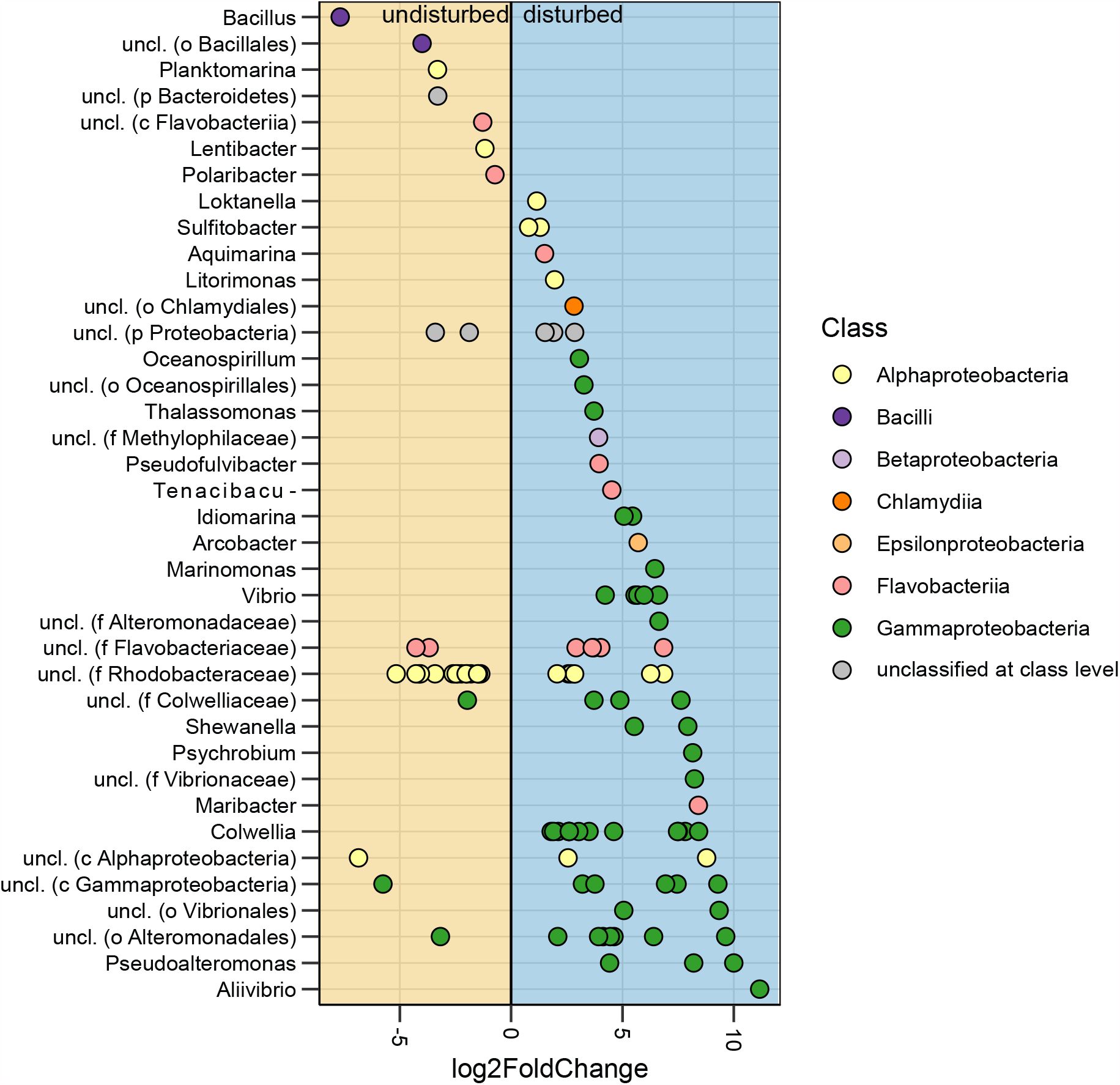
The log2 fold change in relative abundance for the OTUs with a significance level lower than 0.05 (FDR-adjusted DESeq2 p-values). Each point represents an OTU coloured by the class classification. OTUs were grouped according to the genus level. For OTUs that could not be classified at the genus level, the lowest taxonomic classification obtained is indicated in parenthesis (p = phylum, c = class, o = order, f = family, g = genus). OTUs with higher abundance during disturbance are in the blue shaded area, whereas those with higher abundance when the environment was undisturbed are in the orange shaded area.

## Discussion

Predicting community responses to ecosystem changes is essential for improving ecosystem management. From an industrial perspective, we are dependent on stable microbial communities that perform well. Moreover, we live in a time where humans create disturbances at various levels in natural ecosystems. It is therefore important to comprehend the consequences of our activity. To predict the community response to external forces, we need to understand how different ecosystems affect the community assembly processes.

We aimed to fill the knowledge gap on how carrying capacity and periodical disturbances affect the community assembly. It has previously been shown that the carrying capacity affects the community composition (e.g. 44). However, its effect on the assembly processes has remained unclear. Ecosystems with a lower carrying capacity support lower community size. Because the outcome of drift is density-dependent (6), communities with a low carrying capacity should have more populations vulnerable to drifting to extinction. However, our five-times difference in carrying capacity between cultivation regimes did not result in apparent differences in community assembly. The only exception was for the disturbed communities in Period 2, where the low carrying capacity regime (UDL) indicated a stronger influence of selection than the high (UDH; Figure 4b). This observation was surprising as we hypothesised that drift might be more pronounced in systems with lower carrying capacity. In conclusion, the minor effects of carrying capacity observed for the replicate similarity rate for the undisturbed communities suggest that the effect of carrying capacity should be investigated further, including larger differences in carrying capacity.

The effect of the disturbance regime on the microbial community assembly was more evident. The disturbance we investigated was a substantial dilution of the microcosm’s inoculum. The dilution has two significant effects: the community size is reduced, and the concentration of resources increases strongly for the remaining individuals. These two changes are relevant in natural and human-created ecosystems, where resource supply vary due to natural processes (e.g. patchiness and floods) and human activity (e.g. eutrophication and saprobiation).

Investigating the temporal community composition through ordinations can reveal overall successional trajectories (47). We found that whereas the PCoA ordinations indicated an overall deterministic trajectory for the undisturbed communities, the replicate similarity rate indicated that drift dominated the community assembly. This was evident for the microcosms starting with undisturbed culture conditions (UD; Figure 5). However, for the communities going from disturbed to undisturbed conditions (DU), the results were less evident as the replicate similarity rate was around zero. Nonetheless, there was an apparent decrease in the replicate similarity rate when going from disturbed to undisturbed conditions.

The strength and unique feature of our experiment is the crossed design of the disturbance regimes. This crossing considerably increases the robustness of the conclusions drawn from the data. First, during the first period, all microcosms were inoculated with the same community, but in the second period, the twelve communities had assembled individually for 28 days. We could therefore investigate the effects of our experimental variables on drift and selection with different starting conditions. The temporal trends in the data were found to be independent of the starting condition, substantially increasing the strength of our conclusion.

Second, subjecting the communities to the opposite disturbance regime in Period 2 supports that we had stable attractors in our systems. An attractor is a point or a trajectory in the state space of a dynamical system. If the attractor is locally stable, the system will tend to evolve toward it from a wide range of starting conditions and stay close to it even if slightly disturbed (48). We observed locally stable attractors based on the disturbance regime and thus one stationary phase for each disturbance regime. Some ecological systems show dramatic regime shifts between alternative stationary states in response to changes in an external driver (49). Such systems typically exhibit hysteresis in the sense that they will not return directly to the original state by an opposite change in the driver. We found that community composition was reversible and dependent on the disturbance regime, as highlighted by the Bray-Curtis ordinations. This reversibility indicates that the community changes we observed were not catastrophic bifurcations or regime shifts and that it is unlikely that the systems contain multiple stationary states within the same disturbance regime. We think this gives strong support for assuming that drift is the main driver for divergence in the community composition and that selection towards alternative attractors probably plays a minor role. Thus, we can conclude that the shift from a disturbed to an undisturbed ecosystem increased the contribution of drift. Our observations corroborate other investigations of bioreactors (15,50) and simulations (51) that report that stochasticity is fundamental for the assembly of communities. However, the finding that drift was important for structuring the undisturbed microcosms was unexpected.

The selective process competition has been hypothesised to be high in dispersal limited communities where resources are supplied continuously (7), such as in the undisturbed communities examined here. However, our experimental environment offered little variation in the resources provided, as the medium provided was the same throughout the experiment. This may have led to populations becoming “ecologically equivalent”, meaning that their fitness difference was too small to result in competitive exclusion on the time scale of our experiment (5,52). Under these assumptions, community assembly is similar to the neutral model in which the growth rates of the community members are comparable (53).

For the disturbed microcosms, we found that selection dominated community assembly. This finding was also surprising because the disturbance was expected to increase the contribution of drift through relaxation of competition (which stimulates growth of the rare biosphere and strengthens priority effects) and would make low abundance populations vulnerable to local extinction (6,7). During the disturbances, the Sørensen similarity between replicates was stable or increasing, indicating that the periodical disturbance did not result in the extinction of low abundant populations as we hypothesised. Instead, it appears that the dilution removed competition for some time, resulting in a phase where all populations got “a piece of the cake”. Several studies have observed increased stochasticity as a result of increased resource availability (7,11,23,26). However, in our experiment, selection dominated during increased resource availability.

Our results support Zhou et al. 2014 hypothesis stating that determinism should increase due to disturbances in dispersal limited communities (23). However, they contradict their other hypothesis stating that nutrient inputs should increase stochasticity (23). We found that disturbances resulting in periods with exponential growth due to density-independent loss of individuals and high resource input suppressed the effect of stochastic processes. This exponential growth period without competition would enable more populations to stay above the detection limits of the 16S-rDNA-sequencing method.

More OTUs were enriched under the disturbed regime than under the undisturbed. During the disturbance, the microcosms were diluted approximately 2 day^−1^, whereas the dilution factor was 1 day^−1^ during the undisturbed regime. We cannot assume steady state in the disturbed microcosms, but it was interesting to see a substantial increase in the abundance of OTUs classified as *Gammaproteobacteria. Gammaproteobacteria* include many opportunists (54) that appeared to exploit the resource surplus following the disturbance. This opportunistic lifestyle fits within the r- and K-strategist framework (55).

Organisms with high maximum growth rates but low competitive abilities are classified as r-strategists. These r-strategists are superior in environments where the biomass is below the carrying capacity. On the other hand, K-strategists are successful in competitive environments due to their high substrate affinity and resource specialisation (56). Based on the taxonomic responses, it appears as the disturbance selected for r-strategists, whereas the undisturbed regime selected for K-strategists. The r-strategists selected for during the disturbance periods included genera such as *Vibrio* and *Colwellia* (57), and the genus *Vibrio* includes many pathogenic strains (58). Thus, our findings may have implications for land-based aquaculture systems where conditions favouring r-strategists is linked to high mortality and reduced viability of fish (56).

The DeSeq2 results pose some new questions regarding the link between phylogeny and niche fitness. Generally, ecologists assume that closely related taxa have similar niches, as they have a common evolutionary history and, thus, similar physiology (59,60). For example, here, OTUs belonging to *Gammaproteobacteria* co-occurred when the environment was disturbed. However, for other classes such as *Alphaproteobacteria* and *Flavobacteria*, the OTUs responded differently to the disturbance regimes, despite belonging to the same class. This lack of phylogenetically coherent response indicates that the paradigm of correlation between phylogeny and niche requires further studies.

This study was performed on complex marine microbial communities cultivated under controlled experimental conditions. We found that undisturbed environments enhanced the contribution of drift on community assembly and that disturbances increased the effect of selection. These observations might be different in more diverse ecosystems such as soils or the human gut. In such ecosystems, the microbes are more closely associated with, for example, soil particles or attached to the gut lining. It has been shown that the biofilm-associated and planktonic microbial communities have different community composition (61). Consequently, the community assembly processes may be affected differently by environmental fluctuations. Our experimental variables should therefore be tested in other ecosystem settings to verify our conclusions.

To our knowledge, this study is the first to experimentally estimate the effect of periodical disturbances and carrying capacity on community assembly in dispersal-limited ecosystems. We observed that carrying capacity had little effect on community assembly and that undisturbed communities were structured more by drift than disturbed systems dominated by selection. Using an experimental crossover design for the disturbance regime, we showed that these observations were independent of the initial community composition. Our experiment illustrates that cultivating complex natural microbial communities under lab conditions allowed us to test ecologically relevant system variables and draw robust conclusions.

## Acknowledgements

This study was part of the ERA-Net COFASP project “MicStaTech”, funded by the Research Council of Norway (Contract 247558). Financial support was also provided by NTNU, Faculty of Natural Sciences, as a PhD scholarship to MSG and a Fulbright scholarship to IAM. We would like to thank T. Frede Thingstad and the Brendan Bohannan group for valuable comments on an earlier draft of this manuscript.

