## Supplementary figures and methods for "The effect of periodic disturbances and carrying capacity on the significance of selection and drift in complex bacterial communities"

Running title: Effect of periodic disturbance on assembly

### Supplementary figures

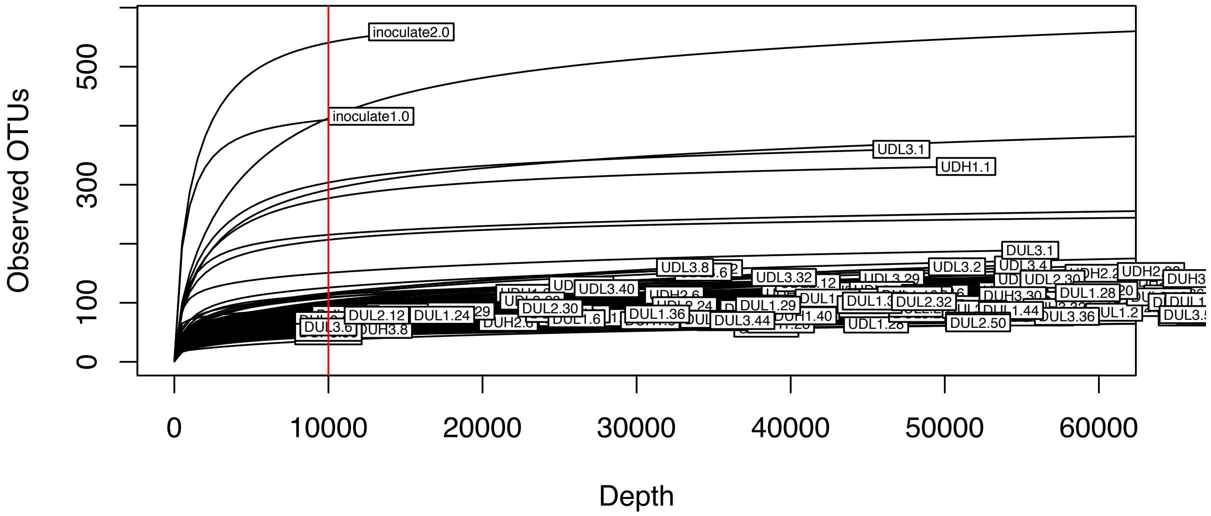


Supplementary figure 1: Rarefaction curves showing the increase in observed OTUs as a function of sequencing depth for each sample. Samples were normalised to 10 000 sequence reads highlighted as the red line. UD were undisturbed the first 28 days and disturbed the last 22 days, while DU was disturbed in the first period and undisturbed in the second. H and L indicates high and low carrying capacity, respectively.

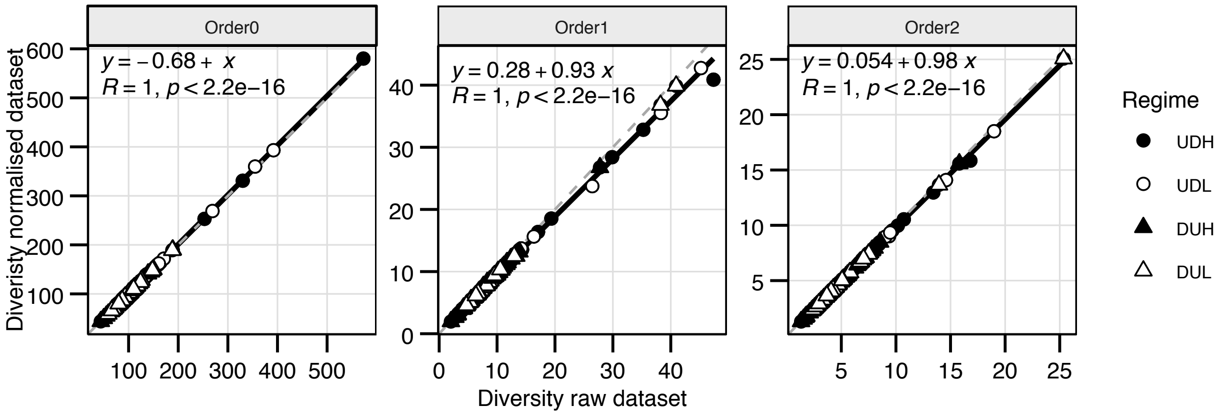


Supplementary figure 2: Relationship between the non-normalised and normalised dataset at the diversity of order 0, 1 and 2. Shapes indicate cultivation regime, formula the linear regression equation and R the Pearson coefficient with accompanying p-value. UD (circles) were undisturbed the first 28 days and disturbed the last 22 days, while DU (triangles) were disturbed in the first period and undisturbed in the second. H (filled) and L (empty) indicates high and low carrying capacity, respectively.

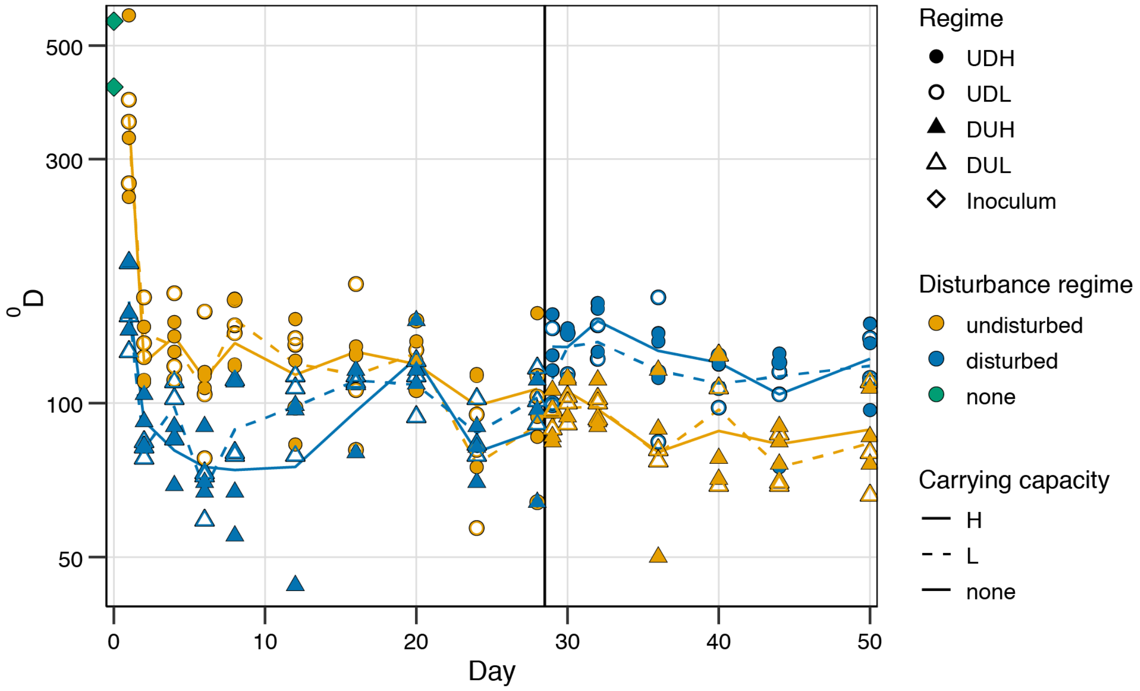


Supplementary figure 3: Richness ($^0$D) as the number of OTUs observed in each sample. Colours indicate the disturbance regime at sampling and shape the overall regime of the sample. UD (circles) were undisturbed the first 28 days and disturbed the last 22 days, while DU (triangles) were disturbed in the first period and undisturbed in the second. H (filled) and L (empty) indicates high and low carrying capacity, respectively. The line is the average richness for each regime at each sampling day (n=3). The inoculum samples (rhombus) were sampled before introduction to lab-culture conditions.

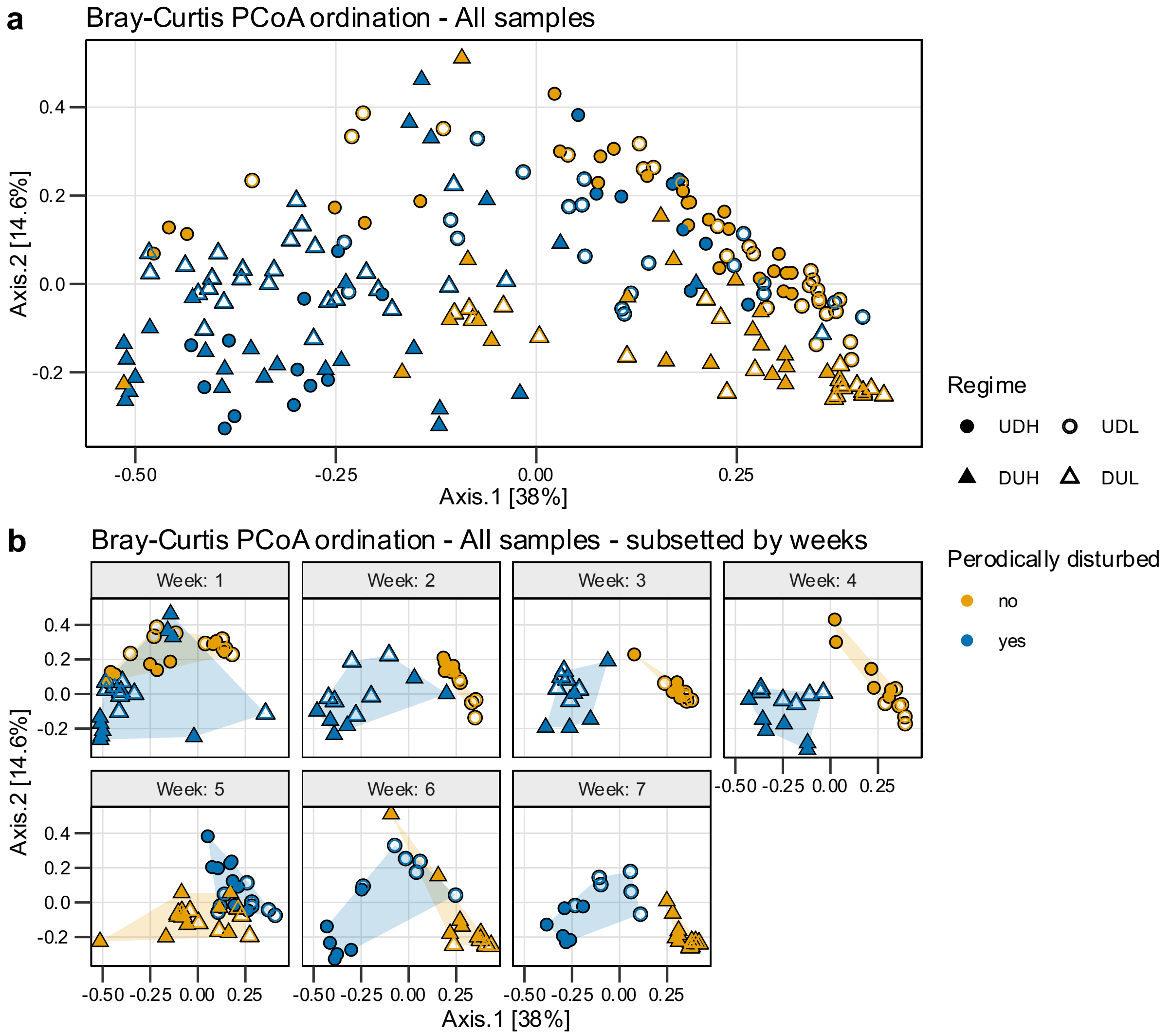


Supplementary figure 4: PCoA ordination based on Bray-Curtis dissimilarity for the bacterial communities. a) The ordination of all samples b) The same ordination space as in a) but subsetted by sampling week to highlight the succession based on the disturbance regime. UD (circles) were undisturbed the first 28 days and disturbed the last 22 days, while DU (triangles) were disturbed in the first period and undisturbed in the second. H (filled) and L (empty) indicates high and low carrying capacity, respectively. Colours represent the disturbance regime at sampling, and the shaded area the spread of samples with similar disturbance regime.

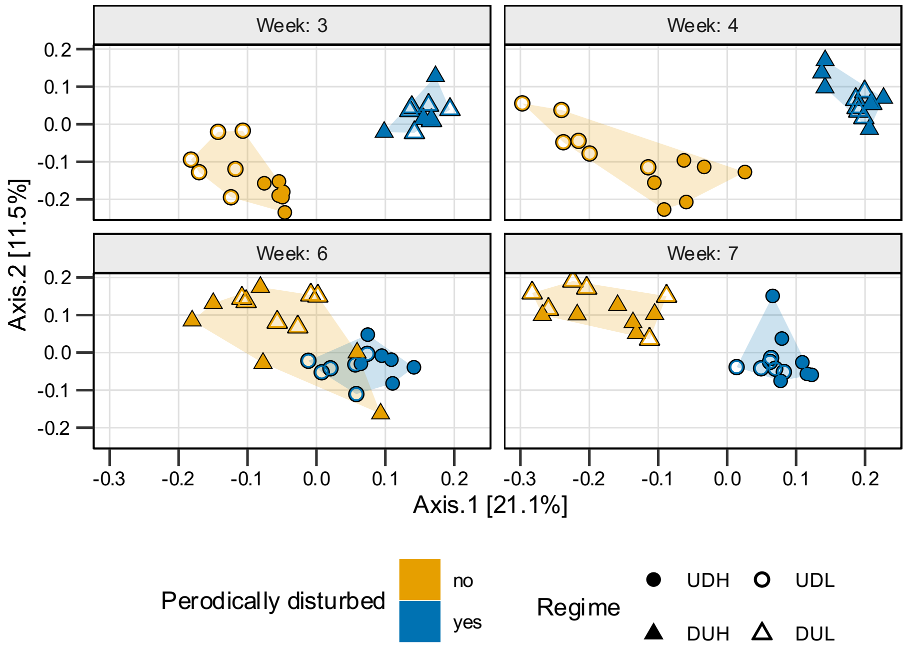


Supplementary figure 5: PCoA ordination based on Sørensen dissimilarity for the bacterial communities at week 3, 4, 6 and 7. The ordination was facetted by week to highlight the succession based on the disturbance regime. UD (circles) were undisturbed the first 28 days and disturbed the last 22 days, while DU (triangles) were disturbed in the first period and undisturbed in the second. H (filled) and L (empty) indicates high and low carrying capacity, respectively. Colours represent the disturbance regime at sampling, and the shaded area the spread of samples with similar disturbance regime.

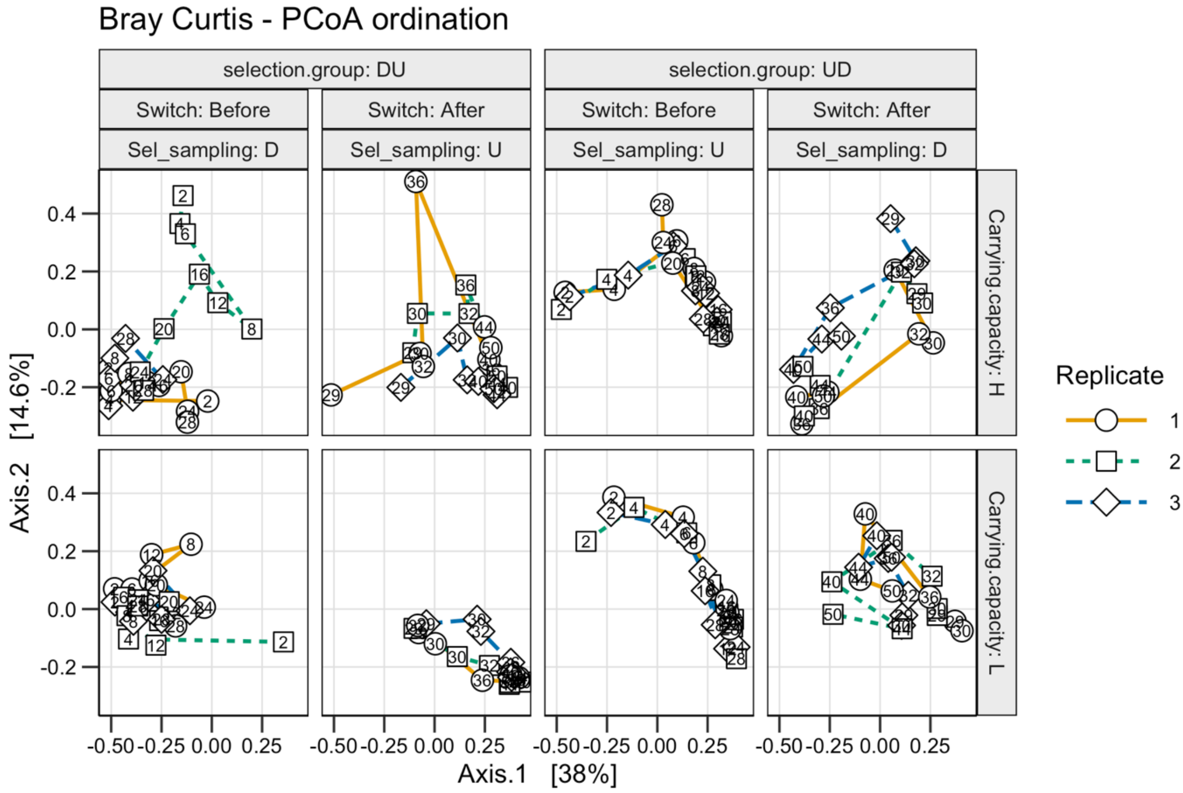


Supplementary figure 6: The temporal trajectories for each replicate microcosms as observed through a PCoA ordination based on Bray-Curtis dissimilarity. The single ordination was split vertically based on carrying capacity and horizontally based on the overall and current disturbance regime. UD were undisturbed the first 28 days and disturbed the last 22 days, while DU were disturbed in the first period and undisturbed in the second. H and L indicates high and low carrying capacity, respectively. Colours and shape differentiate the replicates, with text indicating sampling day.

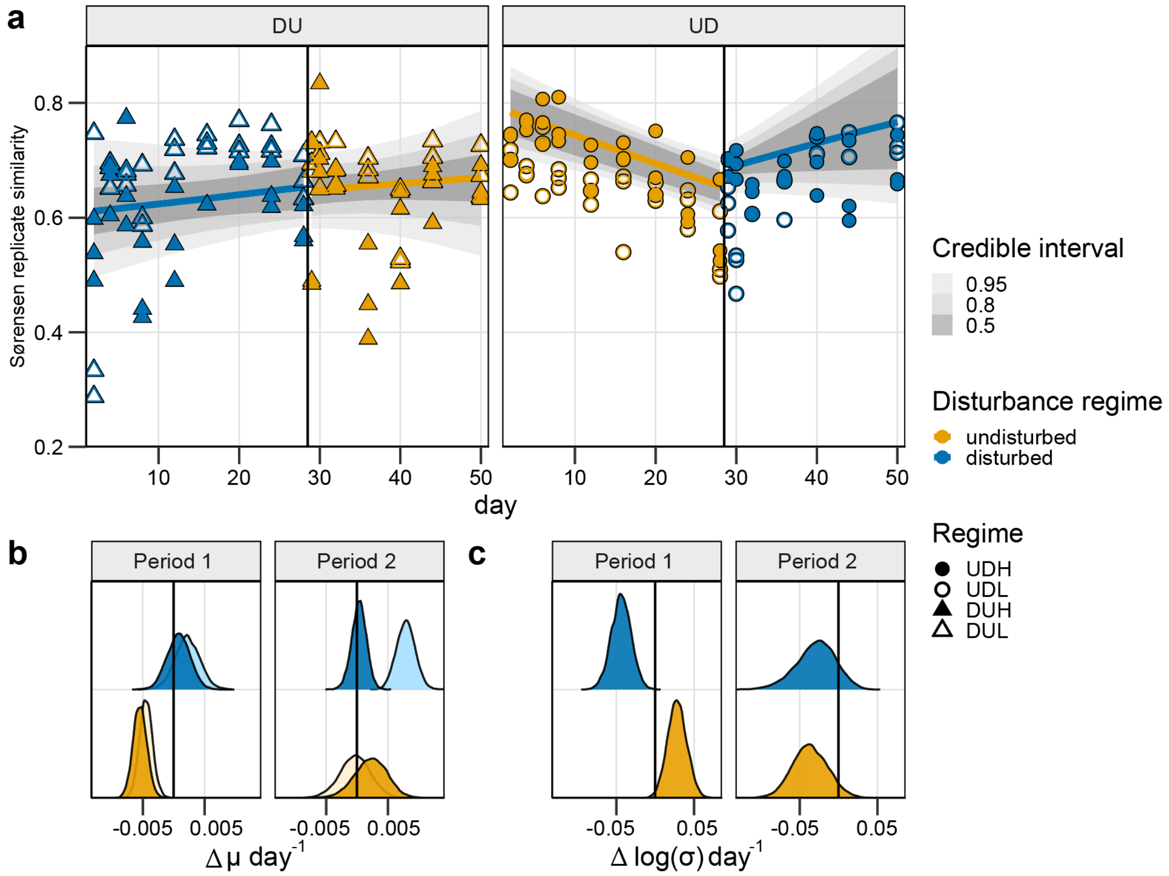


Supplementary figure 7: The Sørensen based models and coefficient estimates for the change in the similarity between replicates over time. **a)** The similarity between replicate communities as a function of time. The models for the replicate similarity change over time are presented as lines with the 0.5, 0,8 and 0.95 credible intervals around it. The observed data used as the response variable in the models are presented as points. UD (circles) were undisturbed the first 28 days and disturbed the last 22 days, while DU (triangles) were disturbed in the first period and undisturbed in the second. H (filled) and L (empty) indicates high and low carrying capacity, respectively. Colours represent the disturbance regime at sampling. **b)** The posterior distributions of the expected replicate similarity ($\mu$) change per day given the interaction between time, the disturbance regime and carrying capacity. The distribution reflects all 8000 estimated replicate similarity changes that would give the observed data. Light and dark colour indicate low and high carrying capacity, respectively. The colour indicates the disturbance regime at sampling. **c)** The posterior distributions for the change in standard deviation per day given the interaction between time and disturbance regime. The distribution reflects all 8000 estimated standard deviation changes per day that would give the observed data. The colour indicates the disturbance regime at sampling.

### Replicate similarity model selection

For each sample, relevant metadata was the disturbance regime at sampling, carrying capacity and sampling day. We calculated the similarity between biological replicates at each sampling day plotted them over time (Supplementary figure 8 a). We also investigated the variation over time in each cultivation regime as the standard deviation. The data indicated a temporal trend regarding both the replicate similarity and the standard deviation as a response to the disturbance regime, especially in Period 1. The replicate similarity appeared to decrease over time in the undisturbed regime and increased when disturbed (Supplementary figure 8 a). The opposite trends were observed for the standard variation (Supplementary figure 8 b).

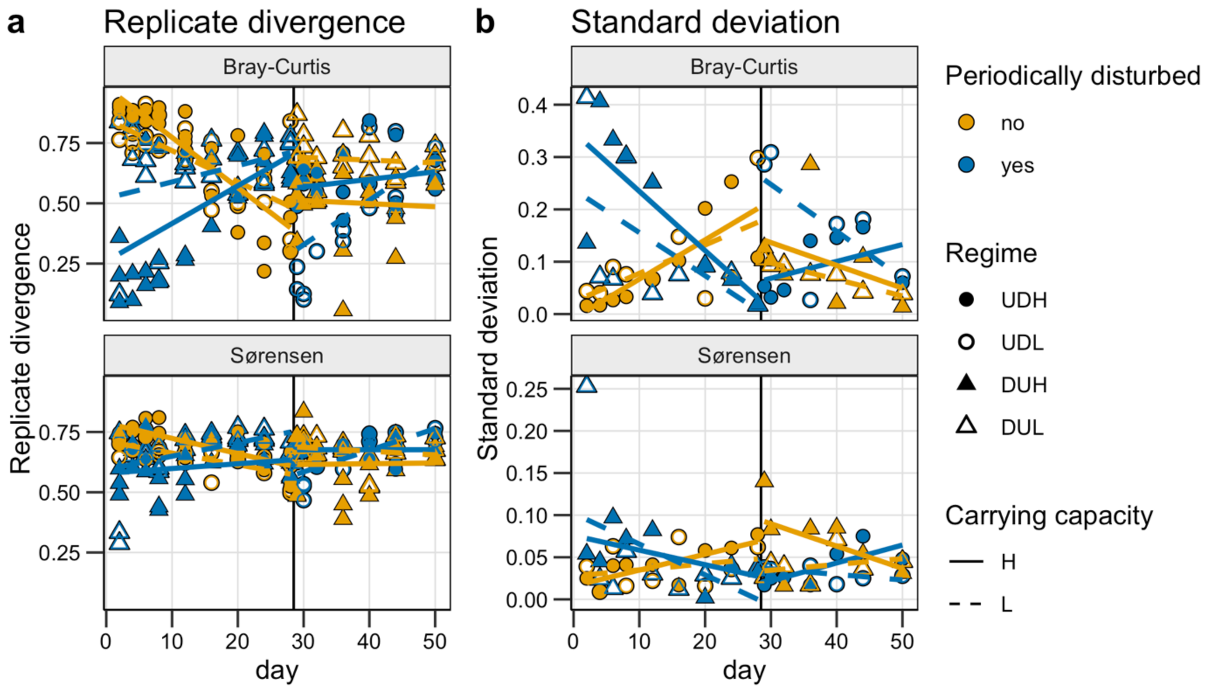


Supplementary figure 8: a) The Bray-Curtis and Sørensen replicate similarity at each sampling day and b) the standard deviation of the similarity at each sampling day. UD (circles) were undisturbed the first 28 days and disturbed the last 22 days, while DU (triangles) were disturbed in the first period and undisturbed in the second. The solid vertical line denotes this switch. H (filled) and L (empty) indicates high and low carrying capacity, respectively. Colours represent the disturbance regime at sampling, and the lines a linear regression within each regime.

Based on these observations, we decided to use a Bayesian hierarchical model approach because this accounts for the limited data points per day, the dependency between sampling days within each microcosm and the heteroscedastic variance (1,2). Bayesian modelling requires that one chooses an appropriate distribution for the data.

We first fitted a normal-, lognormal-, gamma-, beta- and logistic distribution to the data through maximum likelihood estimation for both Bray-Curtis and Sørensen similarity indexes. The function fitdist()  from the package fitdistrplus version 1.0-14 was used for the distribution fitting (3). The observed data were compared with the distributions in a density-, quantile-, cumulative density- and probability plot. Based on these plots, the lognormal, gamma and beta distribution appeared to fit the data well. Because the variation was heteroscedastic, we chose to continue with the normal- and lognormal distributions as these allows constraining the variation to the metadata.

In a first trial, we fitted 24 model-formulas to the temporal Bray-Curtis replicate similarity in Period 1 (Supplementary table 1). The aim was to identify whether lognormal or normal best fitted the data, indicate the model formula, and test whether sampling day should be centralised. Mean-centralising continuous variables can help to reduce covariance between random effect slopes and intercepts (4). For each model, we calculated the predictive density with the leave-one-out-cross validation method through loo(model, reloo = TRUE) from the package loo (version 2.2.0) (5). Then, all the 48 models predictive densities were compared through loo_compare() to identify the model that best fitted the data. The comparison of the predictive densities indicated that the normal distribution was the appropriate distribution, that variance should be given by sampling-day (time) and disturbance regime, and centralising sampling day (time) appeared better (Supplementary table 1).

Supplementary table 1: A total of 48 models were fitted to the Bray-Curtis replicate similarity in Period 1 to get an overview of the appropriate model and sigma formula. The predictive density of each model was estimated, and all models compared to each other yielding the difference in expected log pointwise predictive density(elpd_diff). RS = replicate similarity, comp = replicates compared and S = sigma.

| # | Model formula | Sigma formula | elpd_diff log | elpd_diff nor |
|---|---|---|---|---|
| 1 | RS~ time * disturbed* carrying capacity + (time \| comp) | S ~ 1 | -54.5 | -20.4 |
| 2 | RS~ time * disturbed + (time \| comp) | S ~ 1 | -52.7 | -19.3 |
| 3 | RS~ time * disturbed* carrying capacity + (1 \| comp) | S ~ 1 | -58.1 | -23.8 |
| 4 | RS~ time * disturbed + (1 \| comp) | S ~ 1 | -57.7 | -24.1 |
| 5 | RS~ time * disturbed* carrying capacity + (time \| comp) | S ~ time | -47.8 | -19.1 |
| 6 | RS~ time * disturbed + (time \| comp) | S ~ time | -46.6 | -18.1 |
| 7 | RS~ time * disturbed* carrying capacity + (1 \| comp) | S ~ time | -52.5 | -23.5 |
| 8 | RS~ time * disturbed + (1 \| comp) | S ~ time | -51.9 | -24.5 |
| 9 | RS~ time * disturbed* carrying capacity + (time \| comp) | S ~ time* disturbed | -5.3 | -1.2 |
| 10 | RS~ time * disturbed + (time \| comp) | S ~ time* disturbed | -6.5 | 0.0 |
| 11 | RS~ time * disturbed* carrying capacity + (1 \| comp) | S ~ time* disturbed | -3.3 | -0.7 |
| 12 | RS~ time * disturbed + (1 \| comp) | S ~ time* disturbed | -6.8 | -3.2 |
| 13 | RS~ day * disturbed* carrying capacity + (day \| comp) | S ~ 1 | -53.6 | -20.8 |
| 14 | RS~ day * disturbed + (day \| comp) | S ~ 1 | -52.6 | -19.2 |
| 15 | RS~ day * disturbed* carrying capacity + (1 \| comp) | S ~ 1 | -58.0 | -23.9 |
| 16 | RS~ day * disturbed + (1 \| comp) | S ~ 1 | -57.6 | -24.2 |
| 17 | RS~ day * disturbed* carrying capacity + (day \| comp) | S ~ day | -48.3 | -19.8 |
| 18 | RS~ day * disturbed + (day \| comp) | S ~ day | -46.7 | -18.4 |
| 19 | RS~ day * disturbed* carrying capacity + (1 \| comp) | S ~ day | -52.0 | -23.9 |
| 20 | RS~ day * disturbed + (1 \| comp) | S ~ day | -52.1 | -24.5 |
| 21 | RS~ day * disturbed* carrying capacity + (day \| comp) | S ~ day * disturbed | -4.9 | -0.05 |
| 22 | RS~ day * disturbed + (day \| comp) | S ~ day * disturbed | -7.0 | -0.7 |
| 23 | RS~ day * disturbed* carrying capacity + (1 \| comp) | S ~ day * disturbed | -7.0 | -0.7 |
| 24 | RS~ day * disturbed + (1 \| comp) | S ~ day * disturbed | -6.9 | -3.6 |

As an assurance to continue model selection with the appropriate formulas, we estimated model 1, 5, 9- 13, 17 and 21-24 with the dataset for Bray-Curtis replicate similarity in Period 2 and for Sørensen in both periods. Within each dataset, the models' predictive densities were compared (Supplementary figure 9). Models 21 and 22 were removed due to a high elpd_diff (elpd_diff > 135). Model 23 had the highest predictive density for all the datasets. However, the models with day did not converge fully, and therefore, model 11 with centralised time was chosen.

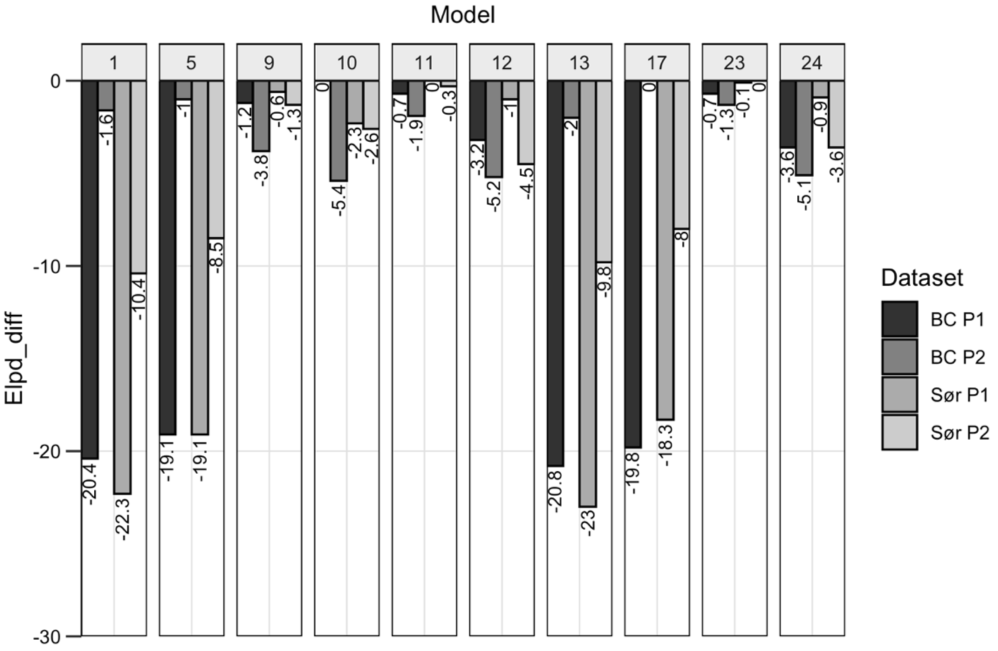


Supplementary figure 9: The difference in the models expected log pointwise predictive density (elpd_diff) within each dataset. A total of 12 models were estimated for each of the datasets to determine if conclusions from the model selection in Period 1 of Bray-Curtis replicate similarity were appropriate for the other datasets. The predictive density of each model was estimated, and all models within the same dataset were compared to each other yielding the difference in expected log pointwise predictive density(elpd_diff). The datasets used were the replicate similarity based on Bray-Curtis in period 1 (BC P1) and 2 (BC P2) and Sørensen in period 1 (Sør P1) and 2 (Sør P2).

We then tested twenty formula variations of model 11 and compared the models' predictive densities (Supplementary table 2). Model 11.1 had the overall highest predictive density and was therefore chosen to estimate the replicate similarity. The estimates based on each similarity index and period are presented in

Supplementary table 3.

Supplementary table 2: A total of 20 models were estimated for the Bray-Curtis (BC) and Sørensen (Sør) based similarity decay in both periods (P1 and P2). The predictive density of each model was estimated, and all models compared to each other yielding the difference in expected log pointwise predictive density(elpd_diff). RS= replicate divergence, comp = replicates compared and S = sigma.

| # | Model formulas | Sigma formulas | Elpd_diff BC P1 | Elpd_diff BC P2 | Elpd_diff Sør P1 | Elpd_diff Sør P2 |
|---|---|---|---|---|---|---|
| **11.1** | RS~ time * disturbed * carrying capacity + (1 \| comp) | S ~time*disturbed | -0.3 | -0.3 | 0.0 | -11.4 |
| **11.2** | | S ~time + disturbed | -17.2 | -17.2 | -9.5 | -10.3 |
| **11.3** | | S ~time*disturbed*carrying capacity | 0.0 | 0.0 | -90.3 | 0.0 |
| **11.4** | | S ~time*disturbed + carrying capacity | -1.2 | -1.2 | -1.4 | -12.6 |
| **11.5** | | S ~time+disturbed + carrying capacity | -18.8 | -18.8 | -10.5 | -11.7 |
| **11.6** | RS~ time * disturbed + carrying capacity + (1 \| comp) | S ~time*disturbed | -3.5 | -3.5 | -1.5 | -14.6 |
| **11.7** | | S ~time + disturbed | -19.2 | -19.2 | -12.3 | -15.4 |
| **11.8** | | S ~time*disturbed*carrying capacity | -2.2 | -2.2 | -14.4 | -3.2 |
| **11.9** | | S ~time*disturbed + carrying capacity | -5.1 | -5.1 | -2.5 | -13.5 |
| **11.10** | | S ~time+disturbed+carrying capacity | -20.6 | -20.6 | -14.1 | -17.4 |
| **11.11** | RS~ time + disturbed + carrying capacity + (1 \| comp) | S ~time*disturbed | -27.3 | -27.3 | -5.8 | -13.8 |
| **11.12** | | S ~time + disturbed | -39.0 | -39.0 | -24.4 | -14.3 |
| **11.13** | | S ~time*disturbed*carrying capacity | -19.1 | -19.1 | -9.6 | -4.0 |
| **11.14** | | S ~time*disturbed + carrying capacity | -28.2 | -28.2 | -6.4 | -14.2 |
| **11.15** | | S ~time+disturbed + carrying capacity | -40.3 | -40.3 | -25.9 | -15.3 |
| **11.16** | RS~ time + disturbed + (1 \| comparison) | S ~time*disturbed | -28.1 | -28.1 | -5.7 | -14.9 |
| **11.17** | | S ~time + disturbed | -38.2 | -38.2 | -24.2 | -15.1 |
| **11.18** | | S ~time*disturbed*carrying capacity | -23.5 | -23.5 | -9.5 | -4.6 |
| **11.19** | | S ~time*disturbed + carrying capacity | -28.9 | -28.9 | -6.0 | -15.4 |
| **11.20** | | S ~time+disturbed + carrying capacity | -39.7 | -39.7 | -25.8 | -16.4 |

Supplementary table 3: Replicate divergence model estimates based on the Bray-Curtis (BC) and Sørensen (Sør) similarity decay in period 1 (P1) and 2 (P2). The model formula used was RS~ time * disturbed * carrying capacity + (1 | comp) and sigma ~time*disturbed.

| | | BC P1 | BC P2 | Sør P1 | Sør P2 |
|---|---|---|---|---|---|
| **Group level** | sd(Intercept) | 0.02 | 0.07 | 0.01 | 0.03 |
| **Population-Level Effects:** | Intercept | 0.72 | 0.50 | 0.71 | 0.61 |
| | sigma_Intercept | -2.41 | -2.10 | -3.15 | -2.46 |
| | time | -0.02 | 0.00 | -0.01 | 0.00 |
| | Disturbed_Yes | -0.24 | 0.10 | -0.09 | 0.06 |
| | Carrying_capacity_L | -0.04 | 0.18 | -0.06 | 0.06 |
| | time:disturbed_Yes | 0.03 | 0.00 | 0.01 | -0.00 |
| | time:Carrying_capacity_L | 0.01 | -0.00 | 0.00 | -0.00 |
| | disturbed_Yes:carrying capacity_L | 0.16 | -0.30 | 0.13 | -0.08 |
| | time:disturbed_Yes:Carrying_capacity_L | -0.01 | 0.02 | 0.00 | 0.01 |
| | sigma_time | 0.06 | -0.03 | 0.03 | -0.04 |
| | sigma_disturbed_Yes | 0.63 | 0.01 | 0.69 | -0.71 |
| | sigma_time:disturbed_Yes | -0.12 | -0.02 | -0.07 | 0.01 |
